# Unique interactions and functions of the mitochondrial small Tims in *Trypanosoma brucei*

**DOI:** 10.1101/2023.05.29.542777

**Authors:** Linda S. Quiñones Guillén, Fidel Soto Gonzalez, Chauncey Darden, Muhammad Khan, Anuj Tripathi, Joseph T. Smith, Ayorinde Cooley, Victor Paromov, Jamaine Davis, Smita Misra, Minu Chaudhuri

## Abstract

*Trypanosoma brucei* is an early divergent parasitic protozoan that causes a fatal disease, African trypanosomiasis. *T. brucei* possesses a unique and essential translocase of the mitochondrial inner membrane, the TbTIM17 complex. TbTim17 associates with 6 small TbTims, (TbTim9, TbTim10, TbTim11, TbTim12, TbTim13, and TbTim8/13). However, the interaction pattern of the small TbTims with each other and TbTim17 are not clear. Here, we demonstrated by yeast two-hybrid (Y2H) analysis that all six small TbTims interact with each other, but stronger interactions were found among TbTim8/13, TbTim9, and TbTim10. Each of the small TbTims also interact directly with the C-terminal region of TbTim17. RNAi studies indicated that among all small TbTims, TbTim13 is most crucial to maintain the steady-state levels of the TbTIM17 complex. Co-immunoprecipitation analyses from *T. brucei* mitochondrial extracts also showed that TbTim10 has a stronger association with TbTim9 and TbTim8/13, but a weaker association with TbTim13, whereas TbTim13 has a stronger connection with TbTim17. Analysis of the small TbTim complexes by size exclusion chromatography revealed that each small TbTim, except TbTim13, is present in ∼70 kDa complexes, which could be heterohexameric forms of the small TbTims. However, TbTim13 is primarily present in the larger complex (>800 kDa) and co-fractionated with TbTim17. Altogether, our results demonstrated that TbTim13 is a part of the TbTIM complex and the smaller complexes of the small TbTims likely interact with the larger complex dynamically. Therefore, relative to other eukaryotes, the architecture and function of the small TbTim complexes are specific in *T. brucei*.

---

African trypanosomiasis is a deadly disease affecting humans and livestock of sub-Saharan African countries (1, 2). The causative agent of this disease is a parasitic protozoan known as *Trypanosoma brucei* (3). In eukaryotes, mitochondria play essential roles in many cellular functions (4–6). This is especially true for *T. brucei,* which only possesses one mitochondrion (7, 8). Mitochondria are home to hundreds of proteins, and the majority (98%) of these proteins are encoded in the nucleus. Thus, this makes the biological process of protein import essential (9). Mitochondrial protein import machinery is composed of several multi-protein complexes, which have been elaborately studied in fungi and mammals (10, 11). We and others identified a non-canonical protein import apparatus in *T. brucei*, which belongs to a group of early divergent eukaryotes (12, 13). Therefore, this unique mitochondrial complex deserves thorough investigation not only to identify novel targets for chemotherapy but also for elucidating the evolutionary perspective of this essential cellular process.

In yeast, mitochondrial protein import machinery is composed of three major complexes: the translocase of the outer membrane (TOM), and two translocases of the inner membrane (TIM), TIM22 and TIM23 (10). The TIM23 complex imports primarily matrix-targeted proteins with the help of a pre-sequence translocase-associated motor (PAM) complex (14, 15). There are a few mitochondrial inner membrane (MIM) proteins imported via TIM23 complex that have a sorting signal and are laterally sorted into the MIM (10). The TIM22 complex import multi-spanning MIM proteins (16, 17). These hydrophobic proteins are chaperoned from the TOM to the TIM22 complex by the help of the small Tim complexes (18–21). As the name implies, these small Tims are smaller proteins, ranging from 8-12 kDa in size. The small Tims are soluble proteins localized in the intermembrane space (IMS) of the mitochondria and these are also associated with the translocase complex. *Saccharomyces cerevisiae* has 5 small Tims: Tim8, Tim9, Tim10, Tim12 and Tim13. The small Tims possess the highly conserved twin CX_3_C motifs. Cysteine residues in these motifs form two intramolecular disulfide bonds. Yeast small Tims form 3 hetero-hexameric complexes of ∼70kDa, (Tim9)_3_-(Tim10)_3_, (Tim9)_3_-(Tim10)_2_-(Tim12), and (Tim8)_3_-(Tim13)_3_ (22, 23). Tim12 associates with the TIM22 translocase and helps to dock the Tim9-Tim10 complex carrying the cargo proteins to the TIM22 complex for further translocation. Humans have Tim8a, Tim8b, Tim9, Tim10a, Tim10b, and Tim13 (24). They also form similar heterohexameric complexes, where human Tim10b acts like Tim12 in yeast (25). In yeast, Tim9, Tim10, and Tim12, are essential, whereas Tim8 and Tim13 are dispensable (18–20). However, in humans, the mutation of Tim8 leads to a neurodegenerative disorder known as the human deafness dystonia syndrome (26). This is likely due to impaired import and assembly of the Tim23 into mitochondrial inner membrane (27). The Tim8-Tim13 complex also plays a role in the assembly of the complex IV in human mitochondria (28).

The crystal structures of yeast and human Tim9-Tim10 complexes and the human Tim8-Tim13 complex revealed that, in each complex, two monomers are alternatively arranged and formed a donut-shaped structure. The core region of this structure consists of the central loop, and the pair of disulfide bonds that form the flat face along the molecular axis, and the N- and C-termini that emanate from the opposite face as tentacle-like structures (18, 29, 30). Both hydrophobic and electrostatic interactions hold the monomers together. Salt bridges formed between highly conserved glutamic acid and lysine residues are found critical for complex stability (24).

*T. brucei* possesses 6 small TbTims (TbTim9, TbTim10, TbTim11, TbTim12, TbTim13, and TbTim8/13). Among these, TbTim9, TbTim10, and TbTim8/13 were initially identified by homology search in *T. brucei* genome data base using Hidden Markov prediction models (32). Tim8/13 showed homology to both Tim8 and Tim13 in other eukaryotes. Later, other small TbTims, TbTim11, TbTim12, and TbTim13, were identified by SILAC-coimmunoprecipitation analysis of TbTim17 and found unique to trypanosomes (33). Except TbTim12, each of these small TbTims possesses the conserved twin CX_3_C motifs. TbTim12 has one Cysteine (C) in each motif, however, alpha-fold prediction showed that all these small TbTims have similar secondary and tertiary structures with its homologues (www.tritrypdb.org). The small TbTims are soluble IMS proteins that are essential for *T. brucei* cell growth and are associated with the TbTIM17 complex (34, 35). In *T. brucei*, TbTIM17 complex imports both the N-terminal- and internal targeting signal-containing nuclear-encoded proteins destined to the mitochondrial matrix or the MIM, respectively (33, 36, 37). Besides TbTim17 and TbTim50, other components of the TbTIM complex are unique to *T. brucei* (34, 35). Previous studies indicate that small TbTims are present in smaller complexes (∼150 kDa and ∼70 kDa) and in a larger complex that is similar in size to the TbTIM17 complex (34, 35). However, it is not clear if these proteins randomly interact with each other and form these complexes or there is a specific pattern and functions of small TbTim complexes in *T. brucei*. Here, we systematically analyzed the interaction pattern and the complexes formed by the small TbTims and compared the effect of RNAi of each individual small TbTim on the integrity of the TbTIM17 complex. Altogether, we found that TbTim9, TbTim8/13, and TbTim10 interact strongly and may form the core small TbTim complex. Except TbTim13, small TbTims form 70 kDa complexes, whereas TbTim13 only associates with the larger complex (>800 kDa) along with TbTim17. TbTim13 is the most crucial among all other small TbTims for TbTIM17 complex biogenesis/stability. Therefore, despite overall structural homology with the small Tims in other eukaryotes, the interaction pattern, and the architecture of the small TbTim complexes are different in *T. brucei*.

## Results

### Small TbTims directly interact with each other with stronger interactions among TbTim9, TbTim10, and TbTim8/13

To determine the interaction pattern of the small TbTims, we used yeast two-hybrid (Y2H) analysis. Each of the six small TbTims was cloned in the pGADT7 and pGBKT7 vectors containing the activation domain (AD) and the DNA-binding domain (BD), respectively, of the GAL4 transcription factor in frame with the insert. The pGADT7 and pGBKT7 plasmids also possess the Myc and HA epitope tag sequences in frame, respectively, to express the corresponding fusion proteins with these tags at the C-terminus. Yeast cells (Y2HGold strain) were co-transfected with the pair of small TbTim clones in all possible combinations and allowed to grow in synthetically defined (SD) double dropout (DDO) plates, lacking leucine (-leu) and tryptophan (-trp). To test the direct interactions among the small TbTims and to differentiate between stronger and weaker interactions, isolated colonies from DDO plates were reinoculated in the triple dropout (TDO) SD plates lacking leucine (-leu), tryptophan (-trp), and histidine (-his), and containing different concentrations (0 mM, 2 mM, 3.5 mM, and 5 mM) of 3-amino-1,2,4-triazole (AT), a histidine synthase inhibitor. At least 3 colonies from each DDO plate were tested for interaction studies. As the negative and positive controls, yeast strain transformed with no DNA and co-transformed with SV40-AD and p53-BD clones, respectively, were plated in parallel in all different TDO plates.

The results showed that yeast cells co-transformed with different pairs of small TbTim clones grew well at 0 mM AT (Fig. 1A-E, plates-I), indicating that the small TbTims were capable to directly interact with each other. However, many of these co-transfectants grew differentially with increasing concentration of AT (Plates II, III, and IV) (Fig. 1A-E, Table-1). Yeast cell co-transformed with the plasmid pairs, TbTim9 with TbTim8/13 (Fig. 1A, pair 7), and TbTim10 with TbTim8/13 (Fig. 1A, pair:8) grew at up to 5.0 mM AT. Whereas TbTim11+TbTim11 (Fig. 1B, pair 5), and TbTim13+TbTim12 (Fig. 1D, pair 4), TbTim13+TbTim13 (Fig. 1E, pair 4) co-transfectants showed moderate growth till at 3.5 mM AT. Lastly, TbTim11+TbTim9 (Fig. 1B, pair 3), TbTim11+TbTim10 (Fig. 1B, pair 4), TbTim11+TbTim13 (Fig. 1C, Pair 3), TbTim8/13+TbTim11 (Fig. 1C, Pair 4), TbTim10+TbTim12 (Fig. 1C, Pair 6), TbTim8/13+TbTim12 (Fig. 1D, Pair 5), and TbTim8/13+TbTim13 (Fig. 1E, pair 5) pairs showed good growth up to at 2.0 mM AT. Therefore, using Y2-H analysis we observed that TbTim8/13 interacts strongly with TbTim9 and TbTim10, (Fig. 1A-F, Table 1), which is also matched with our previous report (35). We also found moderate interactions between TbTim13 with TbTim9 and TbTim10. Overall interaction patterns among small TbTims are shown in a schematic (Fig. 1F).

**Figure 1.**
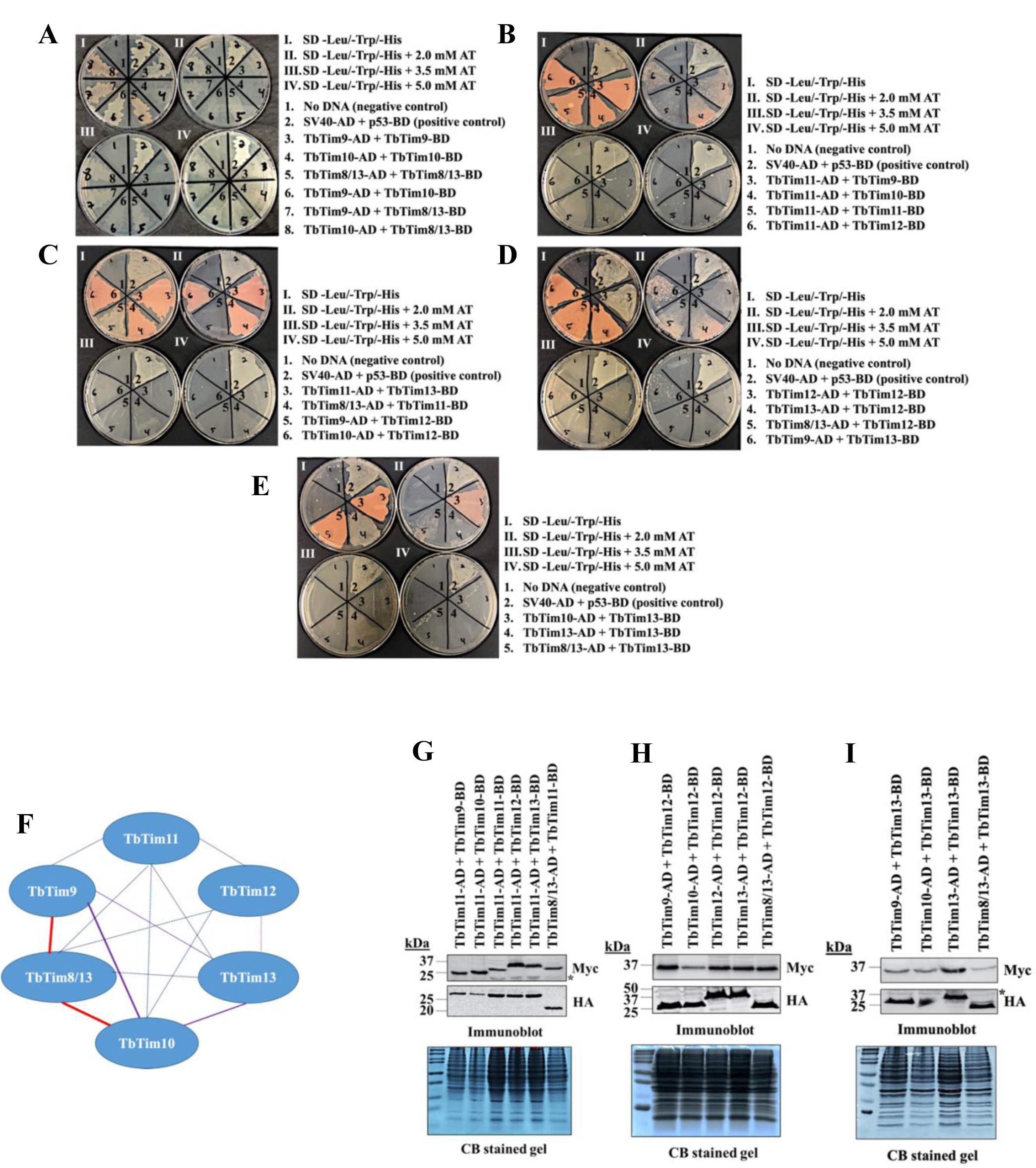
Intermolecular interactions of the small TbTims by Y2H analysis. Six small TbTims were individually cloned in the pGADT7 (Gal4 activation domain, AD) and pGBKT7 (Gal4 DNA-binding domain, BD) plasmids and used as the bait and prey in all possible combinations to investigate their interaction pattern. (A-E)Yeast strain co-transformed with the bait and the prey plasmids were grown in (I) medium lacking leucine, tryptophan, and histidine (–leu/–trp/–his), (II) –leu/–trp/–his medium containing 2.0 mM 3-amino-1,2,4-triazole (AT), (III) –leu/–trp/–his medium containing 3.5 mM AT, and (IV) –leu/–trp/–his medium containing 5.0 mM AT. Yeast cells co-transformed with no DNA was used as negative controls. Yeast cells co-transformed with SV40-AD and p53-BD was used as positive control. The plates shown are representatives of three independent experiments. (F) A schematic of the small TbTim interaction pattern based on the Y2H analysis results. Red lines represent cell growth observed even at 5.0 mM AT, indicating stronger interactions, purple lines represent cell growth observed in the presence up to 2.0 mM AT, indicating a moderate interaction, and dashed blue lines represent cell growth at 0 mM AT, indicating weak interactions. (G-I) Expression of the small TbTims-AD and -BD fusion proteins in yeast. Yeast cells co-transformed with a pair of small TbTims-containing plasmids, as indicated were grown in –Leu/-Trp broth and subjected to protein extraction. Total cellular proteins were analyzed by SDS-PAGE and immunoblot analysis using anti-Myc and anti-HA antibodies to detect BD and AD-fusion proteins tagged with the Myc and HA epitopes, respectively. Corresponding Coomassie-stained gels are shown for loading control.

To verify if the AD and BD fusion proteins with small TbTims were expressed in yeast, we grew co-transformants in SD (-leu,-tryp) liquid medium for 48 h. At this point cells were harvested, and total cellular proteins were analyzed by SDS-PAGE and immunoblot analysis using anti-Myc and anti-HA antibodies. We observed that each of these small TbTims fusion proteins were expressed at the expected size in the corresponding co-transfectant yeast cells (Fig. 1G-I). Duplicate Coomassie-stained gels were used as our loading control. The expression levels of most of the fusion proteins were comparable in all transfectants, indicating that differential interaction pattern is likely due to different affinity among these pairs.

### Both helices of the small TbTims are involved in intermolecular interactions

Structural modeling of the small TbTims using alphafold showed that each of these proteins possesses two anti-parallel α-helices connected by a loop, similar to small Tims in human and fungi (Fig. 2A). It has been shown for yeast Tim9-Tim10 complex that both helices of Tim9 are involved in intermolecular interactions, whereas only the C-terminus of Tim10 interacts with Tim9 and the N-terminus is likely involved in substrate recognition (38). Therefore, we wanted to investigate interaction pattern of individual helices of the small TbTims by Y2H analysis. For this purpose, we sub-cloned the N-terminal (Helix 1, H1) and C-terminal (Helix 2, H2) helices of each small TbTims in the pGADT7 and pGBKT7 vectors and performed Y2H analysis for all possible combination pairs. Results from the pairs that showed interactions are shown (Fig. 2B-G, Table-2). We found stronger interactions among the following pairs: TbTim9H1+TbTim12H1 (Fig. 2B, pair 7), TbTim10H1+TbTim11H1 (Fig. 2C, pair 7), TbTim10H1+TbTim12H1 (Fig. 2D, pair 7), TbTim13H1+TbTim12H2 (Fig. 2E, pair 5), TbTim10H1+TbTim8/13H2 (Fig. 2F, pair 4), TbTim8/13H1+TbTim10H2 (Fig. 2F, pair 5), TbTim8/13H1+TbTim12H2 (Fig. 2G, pair 5), and TbTim12H1+TbTim8/13H2 (Fig. 2G, pair 7). Interestingly, both helices of TbTim10, TbTim8/13, and TbTim12 are involved in intermolecular interaction with several TbTim helices (Fig. 2H). Least number of interactions was for TbTim9, TbTim11, and TbTim13 helices (Fig. 2H, Table-2). Multiple interactions of individual helices suggest that small TbTims dynamically interact with each other and may form multiple complexes. TbTim10, TbTim8/13 and TbTim12 could be present in more complexes than others. However, further studies are needed to understand the composition of different complexes.

**Figure 2.**
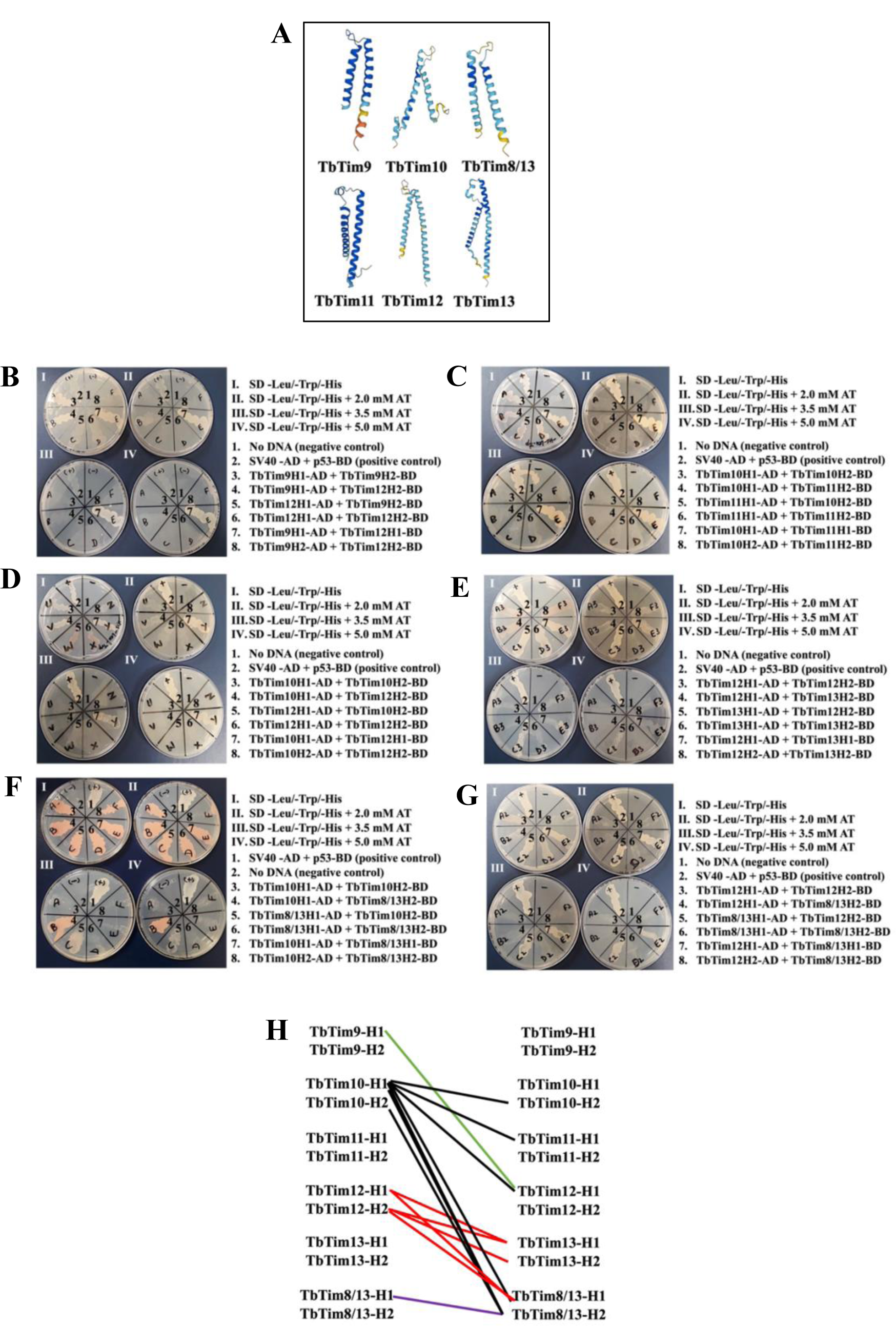
Interaction pattern of individual helices of the small TbTims by Y2H analysis. A) The small TbTims structure models were generated using AlphaFold within the Chimera X software. (B-G) The N-terminal (Helix-1) and the C-terminal (Helix-2) of each of the small TbTims were cloned into pGADT7 and pGBKT7 plasmids. Yeast strain (Gold) were co-transformed with a pair of these plasmids in all possible combinations and grown in (I) –leu/–trp/– his, (II) –leu/–trp/–his medium containing 2.0 mM AT, (III) –leu/–trp/–his medium containing 3.5 mM AT, and (IV) –leu/–trp/–his medium containing 5.0 mM AT. Yeast cells co-transformed with no DNA was used as negative controls. Yeast cells co-transformed with SV40-AD and p53-BD was used as positive control. The plates shown are representatives of three independent experiments. (H) A schematic of the interaction pattern of the small TbTims helices. Only the stronger interacting pairs (growth at 5.0 mM AT) are shown. Interactions of TbTim9, TbTim10, TbTim12, and TbTim8/13 helices with others are shown by green, black, red, and purple lines, respectively.

### TbTim17 C-terminal domain directly interacts with most of the small TbTims

It has been shown that the small TbTims are associated with the TbTIM17 complex (34, 35). However, it is not known if they directly interact with TbTim17. If so, it is unknown which regions of TbTim17 interact with the small TbTims. Like Tim17/22/23 family proteins, TbTim17 possesses 4 predicted transmembrane domains (TMDs) (TMPred, a transmembrane prediction server), whereas the N-terminal (1-30 AAs) and C-terminal (140-152 AAs) hydrophilic regions are exposed in the IMS along with the loop-2 (93-102 AAs), the connecting region between the TMD2 and TMD3 (Fig. 3A). Swiss model structure of TbTim17 based on the cryo-EM structure of the human Tim22 are shown in Fig. 3B. The N- and C-terminals are shown in blue and red, respectively (Fig. 3B). Since small TbTims are localized in the IMS (34, 35), we selected these regions of TbTim17 for testing their interactions with small TbTims by Y2H analysis. We sub-cloned the N- and C-terminal hydrophilic regions, and the loop-2 region of TbTim17 in pGADT7 and pGBKT7 vectors and used these with the AD- and BD-clones for small TbTims for Y2H analysis (Fig. 3C-F, Table 3). Full length TbTim17 was also cloned in the pGADT7 vector and used for Y2H analysis in parallel (Fig. 3F). The co-transformed yeast cell with the N-terminal region of TbTim17 clone and the individual small TbTim clones grew well up to at 2.0 mM AT, but the growth was reduced at 3.5 mM AT, and significantly hampered at 5.0 mM AT (Fig. 3C). Therefore N-terminal region of TbTim17 showed a moderate interaction with the small TbTims. Yeast cells transformed with the TbTim17 loop-2 clone and individual small TbTim clones hardly grew even at 0 mM AT (Fig. 3D). Only a mild growth was observed for the yeast cell transformed with the TbTim17 loop-2 and TbTim11 plasmids. Therefore, we conclude that the loop-2 region of TbTim17 does not interact directly with the small TbTims. Results showed that TbTim17 C-terminal region strongly interacts with all the small TbTims, as the co-transformants were able to grow well up to at 5 mM AT (Fig. 3E, Table-3). We also examined direct interaction between the full length TbTim17 with the small TbTims-BD (Fig. 3F). However, none of these small TbTims showed interactions with the full-length TbTim17 except for TbTim17+TbTim8/13 and TbTim17+TbTim12 pairs, which grew at 0 mM AT only (Fig. 3F, Table-3). The lack of interaction with full length TbTim17 is probably because it is a hydrophobic membrane protein and thus didn’t fold properly when expressed in the yeast cytosol or when it enters the nucleus. Recent cryo-EM data for human TIM22 complex showed that Tim9-Tim10 heterohexameric complexes interact primarily with the N-terminal of Tim22 (39). Our results indicated that in *T. brucei* the C-terminus of TbTim17 is likely holding the small TbTims.

**Figure 3.**
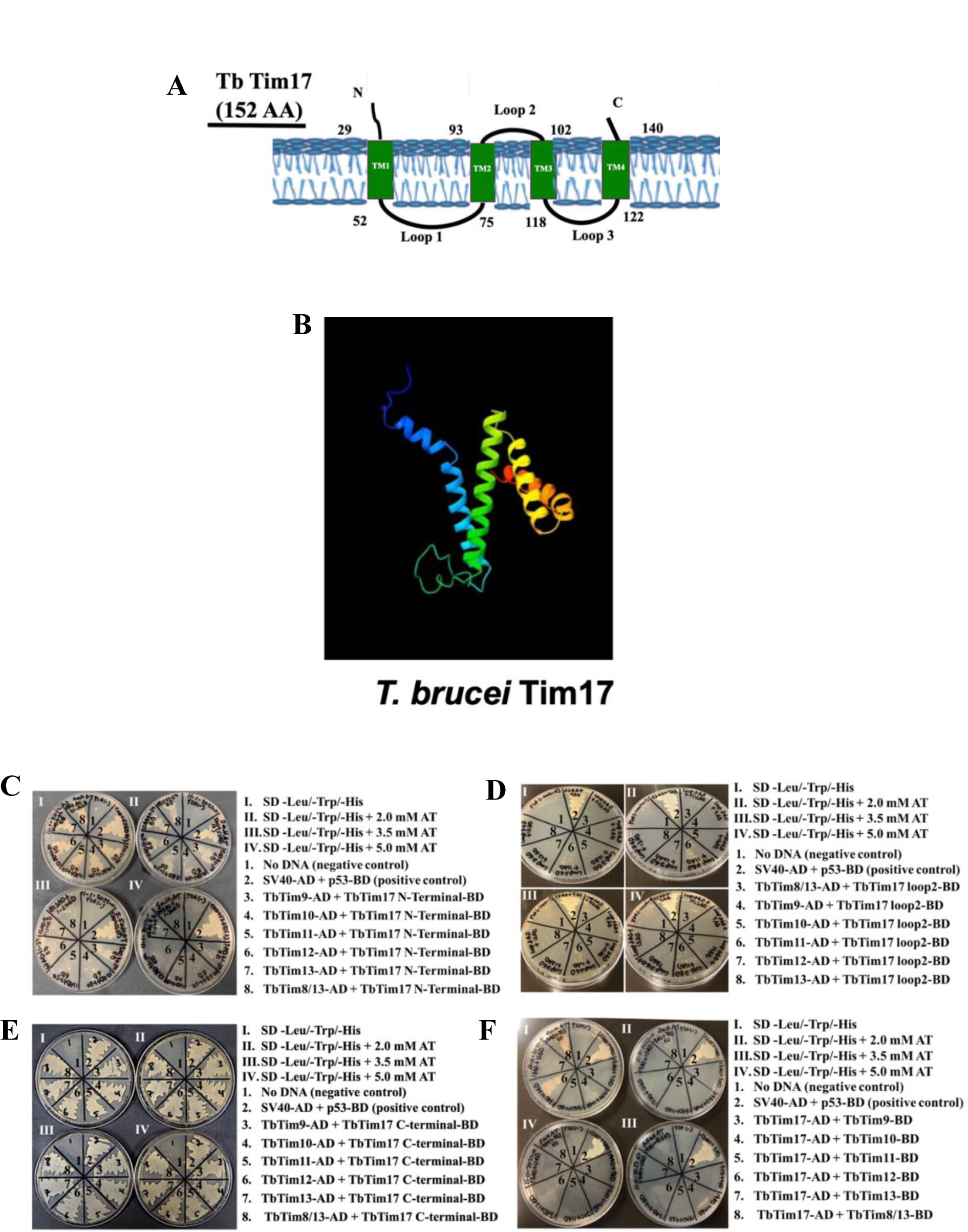
Y2H analysis of the small TbTims interactions with TbTim17. (A) A schematic of TbTim17 protein. The predicted transmembrane domains (TM1-TM4) and the membrane topology are shown. Numbers indicate the amino acid residues on the protein. (B) A structural homology model of TbTim17 based on the cryo EM structure of HsTim22 was generated using Swiss Model prediction software within Chimera X. (C-F) The N-terminal (1-30 AAs), loop 2 (93-102 AAs), and the C-terminal (140-152 AAs) of TbTim17 were sub-cloned in pGBKT7 to use as the bait. The small TbTims cloned in the pGADT7 were used as the prey to investigate interactions between each of the TbTim17 fragments with individual small TbTims. The full legnth (FL) TbTim17 cloned in the pGADT7 and the small TbTims cloned in pGBKT7, were also used in parallel for Y2H analysis. Yeast strain (Gold) co-transformed with the bait and prey plasmids were grown in (I) –leu/–trp/–his, (II) –leu/–trp/–his medium containing 2.0 mM AT, (III) –leu/–trp/–his medium containing 3.5 mM AT, and (IV) –leu/–trp/–his medium containing 5.0 mM AT. Yeast cells co-transformed with no DNA was used as negative controls. Yeast cells co-transformed with SV40-AD and p53-BD was used as positive control. The plates shown are representatives of three independent experiments.

### Among all small TbTims, TbTim13 is most essential for TbTim17 complex stability

We have previously reported that knockdown of TbTim9, TbTim10, TbTim8/13 reduced the levels of TbTim17 and destabilized the TbTIM17 complex (35). Wanger et al., also reported that the TbTim11, TbTim12, and TbTim13 are required for TbTIM17 complex stability (34). To assess if knockdown of any of these small TbTims has a stronger effect than others, we compared the levels of TbTim17 and TbTIM17 complex in mitochondria isolated from each of these six small TbTims knockdown *T. brucei*. For this purpose, we generated TbTim11, TbTim12, and TbTim13 RNA interference (RNAi) cell lines in the same way we previously generated TbTim9, TbTim10, and TbTim8/13 RNAi cell lines. Quantitative RT-PCR analyses showed that TbTim11, TbTim12, and TbTim13 RNAi reduced the levels of the corresponding target transcript about 90%, 60%, and 90%, respectively, by 48 h post-induction (Fig. S1A). We observed a significant growth inhibition of *T. brucei* due to knockdown of TbTim11 or TbTim12 or TbTim13, specifically at day 4 post-induction of RNAi and after (Fig. S1B). Next, each cell line was induced with doxycycline (1.0 μg/ml), cells were harvested at day 2 and 4 post-induction and crude mitochondria were isolated. Analysis of the mitochondrial proteins by SDS-PAGE and immunoblotting using anti-TbTim17 antibody revealed that at day 2 post-induction, TbTim17 levels were moderately reduced in all small TbTim RNAi samples; however, maximum reduction (∼40%) was observed for TbTim13 RNAi (Fig. 4A and B). This trend was continued at day 4 post-induction as we observed that TbTim17 levels were reduced below 10% of the parental control (Fig. 4A and B) in TbTim13 knockdown mitochondria. At this time point TbTim9, TbTim8/13 and TbTim10 knockdown also reduced the TbTim17 levels about 80% and knockdown of TbTim11, and TbTim12 reduced the levels of TbTim17 to 60%. In contrast to TbTim17, RNAi for individual small TbTims affected mHsp70 and VDAC levels similarly. In comparison to control, mHsp70 levels increased due to knockdown of small TbTims, which could be a stress response phenomenon, since the small TbTims are essential for the *T. brucei* procyclic form. Small TbTims knockdown reduced VDAC levels as we reported earlier, suggesting a role of the small TbTims in VDAC biogenesis (Fig. 4A). Tubulin was used as a loading control (Fig. 4A).

**Figure 4.**
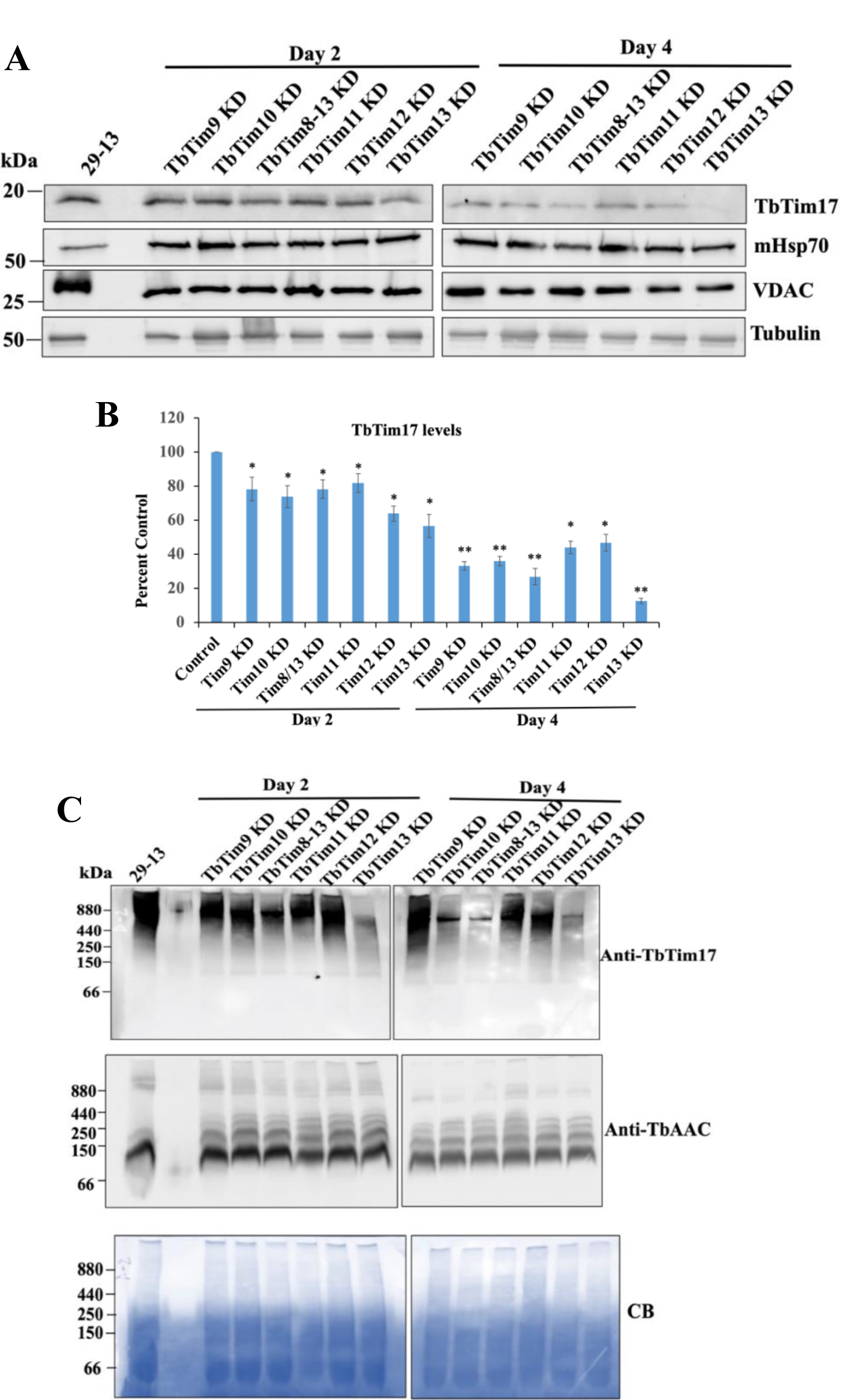
Effect of the knockdown of the small TbTims on TbTim17. (A) The steady-state levels of TbTim17 in the small TbTim RNAi mitochondria. The small TbTim RNAi cells were induced with doxycycline, cells were harvested at day-2 and day-4 after induction for isolation of the mitochondrial fraction. Mitochondrial proteins from the parental (29–13), TbTim9 RNAi (TbTim9 KD), TbTim10 RNAi (TbTim10 KD), TbTim8/13 RNAi (TbTim8/13 KD), TbTim11 RNAi (TbTim11 KD), TbTim12 RNAi (TbTim12 KD) and TbTim13 RNAi (TbTim13 KD) *T. brucei* were analyzed by immunoblot using TbTim17, Hsp70, VDAC, and β-tubulin antibodies. (B) Quantitation of the TbTim17 levels in each RNAi cells. Densitometric scanning of the TbTim17 and tubulin protein bands were performed using ImageJ software. Band intensities for TbTim17 were normalized with the corresponding tubulin bands and calculated relative to the levels of TbTim17 in the 29-13 mitochondria sample. Values were plotted for each RNAi cell lines. Standard errors were calculated from three independent experiments. **, P <0.05. (C) Analysis of the TbTIM17 complex levels in small TbTim knockdown mitochondria. Mitochondria were isolated from the parental (29–13) and the small TbTim RNAi cells grown in the presence of doxycycline for 2 and 4 days. Equal micrograms of mitochondrial proteins were solubilized with digitonin (1%) and analyzed by BN-PAGE followed by immunoblot analysis using anti-TbTim17 and anti-TbAAC antibodies. The position of the molecular weight marker proteins on the gels are shown. Coomassie Blue (CB)-stained gels show equal loading of samples.

Next, we compared the levels of TbTIM17 complex in small TbTim RNAi mitochondria by Blue-Native (BN)-PAGE followed by immunoblot analysis, using TbTim17 antibody. It was noticed that similar to the steady state levels of TbTim17 protein, TbTIM17 complex levels were reduced significantly due to TbTim13 knockdown at day 2 and more at day 4 post-induction (Fig. 4C). We noticed TbTim10 and TbTim8/13 knockdown also significantly reduced TbTIM17 complex levels relatively more compared to TbTim9, TbTim11, and TbTim12 knockdown at day 4 post-induction (Fig. 4C). All these results suggest that TbTim13 is most crucial for the stability or biogenesis of the TbTIM17 complex. TbTim10 and TbTim8/13 also plays significant role for this process and other small TbTims may be secondarily involved in maintaining TbTIM17 complex integrity. Probing the blot with antibody for ADP/ATP carrier (AAC), an abundant MCP, we observed that each of these small TbTims knockdown reduced the levels of AAC matured complex (∼120 kDa) similarly at day 4 post-induction; however, minimal effect was observed at day 2 post-induction, which could be due to a high abundance of this protein in mitochondria. Therefore, at this moment it is difficult to assess if any individual small TbTim has a preferential effect on AAC import. Coomassie stained gel was used as our loading control. Overall, by comparing the effect of RNAi all along we observed that six small TbTims have a differential effect on TbTim17 complex integrity, suggesting a specific role of the individual small TbTim in this process.

### Effect of small TbTim knockdown on mitochondrial proteomes

To understand the effect of small TbTims knockdown on mitochondrial proteomes, we performed semi-quantitative proteomics analysis of the crude mitochondria samples from each small TbTim RNAi cell line and compared that with the parental control (Pro 29-13) mitochondria. Results from three independent biological replicates showed ∼63 proteins upregulated (1.5-to 50-fold) and about 82 proteins down regulated (0.1-to 0.8-fold) with statistical significance (high-moderate) in all RNAi samples (Supplementary File S1, Fig. 5A). A heatmap analysis and interactome studies revealed that a cluster of heat shock proteins and chaperones such as Hsp70, Hsp60, Hsp84, Hsp78 or Clp protease, co-chaperone GrpE, DnaJ50, and glucose regulated protein were upregulated due to knockdown of any of these small TbTims, albeit at different levels (Supplementary File S1). Furthermore, several TCA cycle enzymes that were also upregulated. These included 2-oxogluterate dehydrogenase, dihydrolipoyl dehydrogenase, pyruvate dehydrogenase, and others. Interestingly, the receptor subunit of the ATOM complex, Atom46, and a subunit of the TbTIM17 complex, TbTim42, were down regulated about 5-to 6-fold, in different small TbTims knockdown mitochondria (Supplementary File S1, Fig. 5A). It is possible that downregulation of Atom46 was to reduce the protein load in the import complex as a compensatory mechanism to prevent protein misfolding due to the absence of the small TbTims. TbTim42 levels were reduced, which could be due to a reduction of the levels of the TbTIM17 complex. Among the downregulated proteins we found a few mitochondrial carrier proteins (MCPs), several cytochrome oxidase subunits, tryparedoxin peroxidase, and some glycosomal enzymes like glycerol kinase, glycerol-3-phosphate dehydrogenase, malic enzyme and others (Supplementary File S1, Fig. 5A). The mitochondrial outer membrane (MOM) proteins, Atom40 and POMP10 were downregulated specifically in TbTim10 knockdown mitochondria (Supplementary File S1, Fig. 5A). Interactome studies using STRING program revealed that the upregulated and downregulated proteins were clustered primarily into 2 and 3 groups, respectively (Fig. 5B and C). The two clusters of the upregulated proteins were: 1) chaperones and proteases, and 2) the TCA cycle enzymes (Fig. 5B). Whereas the three clusters of the downregulated proteins include: 1) proteins required for import, folding and redox balance, 2) glycolytic enzymes, and other glycosomal proteins, and 3) a few cytoskeleton-associated proteins (Fig. 5C). Together, these results suggests that small TbTims knockdown generated oxidative stress, likely due to downregulation of mitochondrial energy production due to inhibition of mitochondrial protein import that upregulated chaperone proteins and altered the metabolic pattern in the procyclic form of *T. brucei*.

**Figure 5.**
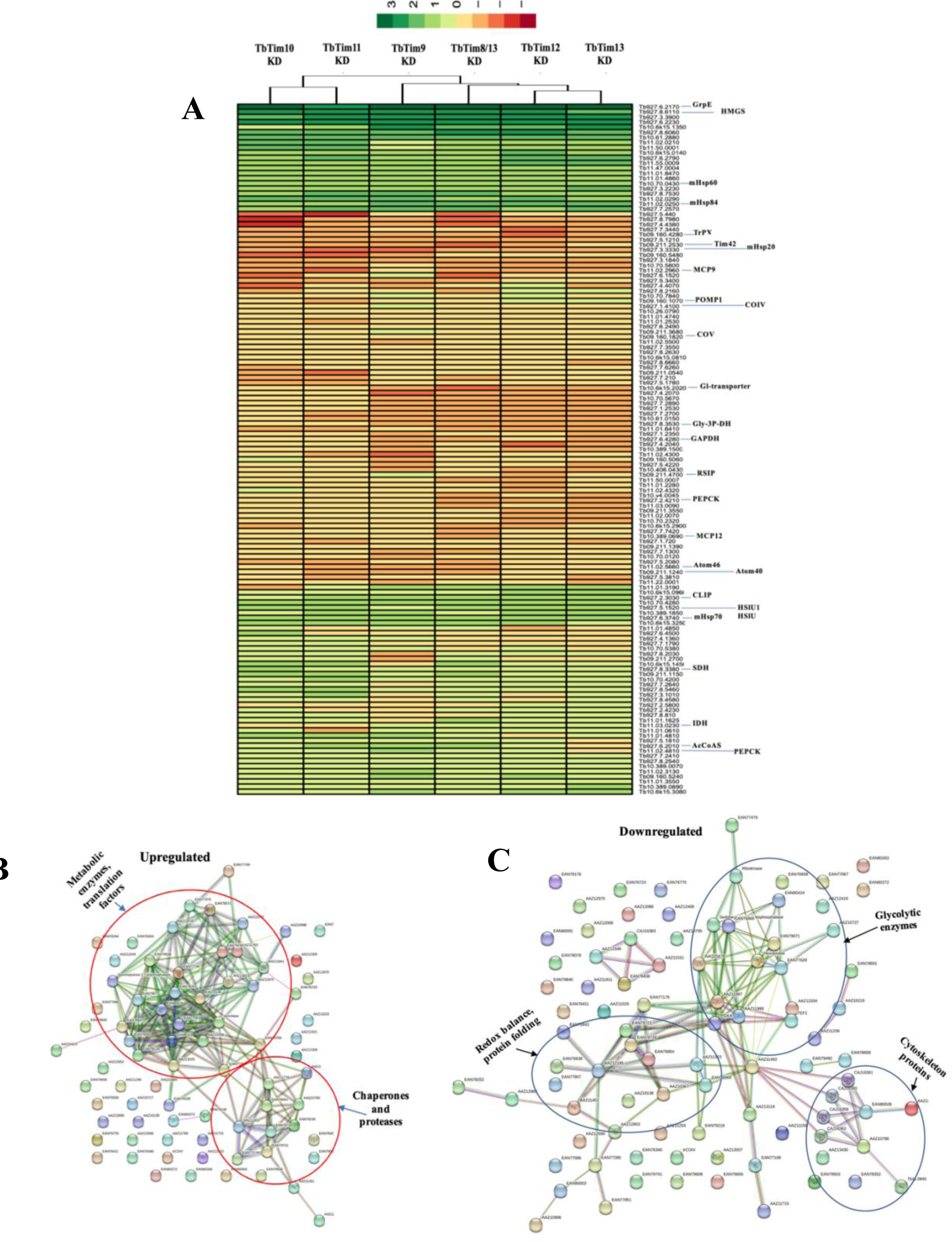
Effects of the small TbTim RNAi on mitochondrial proteomes. The semiquantitative proteomics analysis were performed to compare the proteomes in the parental control and small TbTims knockdown mitochondria. (A) Statistically significant up and down regulated proteins in small TbTims RNAi samples in comparison to control from three biological replicates were used to generate a heatmap using Euclidean distance measure and complete linkage analysis. Each column represents relative levels of proteins in indicated knockdown sample, and each row represents one protein identified within the data set. Visualization was done with the heatmap R package. (B and C) Interaction map of the up and downregulated proteins were used for STRING analysis, as described in the experimental procedure. Clusters of proteins are circled and labeled according to the gene ontology (GO) term.

### Co-immunoprecipitation analyses from *T. brucei* mitochondrial extract showed that TbTim10 has stronger association with TbTim9 and TbTim8/13, and weaker for TbTim13

From our Y2H data we found that small TbTims have differential interaction patterns with each other. To test if that is also true *in T. brucei* mitochondria we performed co-immunoprecipitation analysis of the small TbTims from the *T. brucei* mitochondrial lysate. Due to a lack of availability of good antibodies for small TbTims, we expressed C-terminally tagged small TbTims in pair in *T. brucei*. For this purpose, TbTim10 was tagged at the C-terminus with 2X-Myc and was co-expressed (from an inducible expression vector) in *T. brucei* where either TbTim9, TbTim8/13, TbTim12, or TbTim13 tagged with a C-terminal 3X-HA were expressed (from a constitutive expression vector). We measured the growth kinetics of these cells after induction with doxycycline to see if overexpression of small TbTims has any effect on the cells. Expression of TbTim10-Myc with TbTim9-HA, TbTim10-HA, or TbTim8/13-HA did not show any difference in cell growth (Fig. S2); however, expression of TbTim10-Myc with TbTim12-HA or TbTim13 HA showed inhibition of cell growth, particularly after day 2 post-induction. Therefore, we kept the induction period only 2 days for all cell lines for the subsequent experiments. Next, we checked the expression levels and mitochondrial localization of the tagged small TbTims by immunoblot analysis of the sub-cellular fractions. Both Myc and HA-tagged small TbTims were localized in mitochondria and their expression levels were comparable, except we noticed that TbTim10-Myc levels were reduced when co-expressed with TbTim12-HA or TbTim13-HA (Fig. 6A). Next, we isolated mitochondrial fraction from these cells at day 2 post-induction and mitochondrial proteins were solubilized with 1% digitonin. We have shown that at this condition, small TbTims were still associated with the larger TbTIM17 complexes, and a portion was present in smaller complexes (70 - 150 kDa) by BN-PAGE analysis (35). Mitochondrial extracts were used for immunoprecipitation with either anti-Myc or anti-HA coupled agarose beads and analyzed by immunoblot analysis probing with Myc and HA antibodies. TbTim17 and VDAC were used as positive and negative controls, respectively. Results show that TbTim10-Myc was precipitated by Myc antibodies from all samples, as expected (Fig. 6B, upper panel). This antibody also pulled down TbTim9-HA and TbTim8/13 HA ∼35% and 45% of the input, respectively; however precipitated TbTim10-HA and TbTim12-HA ∼30% and ∼25% of the input, respectively (Fig. 6B and C). In contrast, TbTim13-HA was not co-precipitated with TbTim10-Myc by anti-Myc antibodies (Fig. 6B and C), indicating that TbTim10 has less association with TbTim13. We also observed that TbTim10-Myc co-precipitated a significant fraction of TbTim17 from all samples except when TbTim13-HA was co-expressed with TbTim10-Myc (Fig. 6B), indicating that overexpression of TbTim13 hampered the interaction between TbTim10 with TbTim17. As expected, VDAC was not co-precipitated with TbTim10-Myc. No protein band was detected by HA antibody from samples that only expressed TbTim10-Myc.

**Figure 6.**
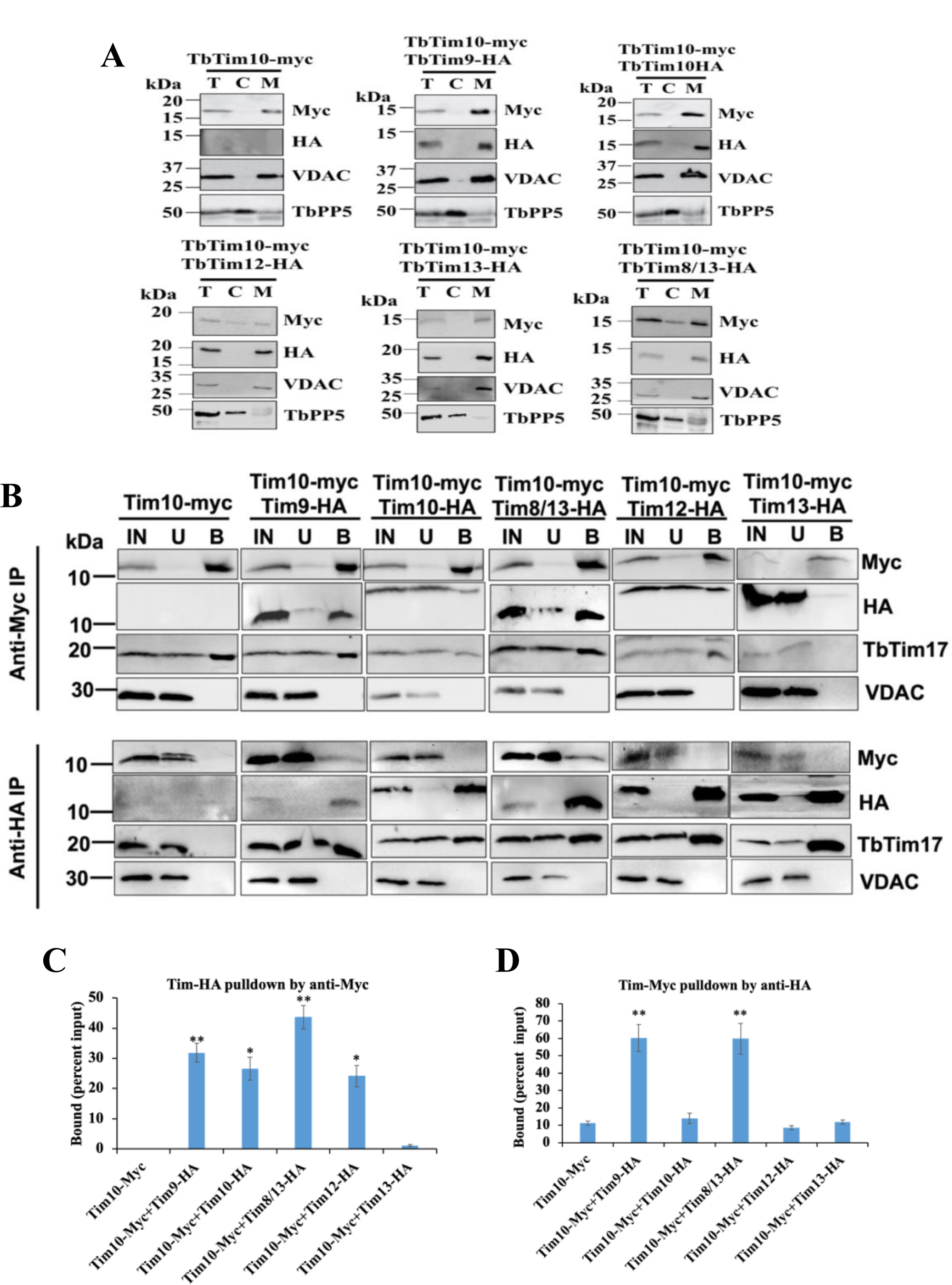
Co-immunoprecipitation analysis of the small TbTims from *T. brucei* mitochondrial extract. *T. brucei* cell line that expresses TbTim10-2X-Myc in a tetracycline inducible fashion were transfected individually with TbTim9-3X-HA, TbTim10-3X-HA, TbTim8/13-3X-HA, TbTim12-3X-HA, TbTim13-3X-HA constitutively expression constructs to generate double-tagged small TbTims cell lines. (A) Localization of the tagged small TbTims. Each cell line was induced for 2 days with doxycycline and subcellular fractions were collected. Proteins from the total cell lysates (T), cytosol (C), and mitochondrial fractions (M) were subjected to SDS-PAGE and were probed with anti-Myc and anti-HA antibody. VDAC and TbPP5 were used as the mitochondrial and cytosolic marker proteins, respectively. (B) Co-immunoprecipitation analysis. The digitonin-solubilized mitochondrial extracts from double-tagged small TbTims cells were used for immunoprecipitation using agarose beads coupled with the anti-Myc or anti-HA antibody. Proteins in the input (IN), unbound (U), and bound (B) fractions were analyzed on SDS-PAGE followed by immunoblot analysis using Myc and HA antibodies. Ten percent (vol/vol) of the IN and U, and 25% (vol/vol) of the B were loaded in the respective lanes. Blots were probed also with TbTim17 and VDAC antibodies. (C and D) Densitometry analysis of the immunoblots using ImageJ software. The quantitated bound fractions (Myc and HA) were calculated as percentage of total input and plotted for each sample. Standard errors were calculated from three independent experiments. Significance values are calculated from t test and are indicated by asterisks as follows; **, p< 0.01; *, p <0.05.

Similarly, when immunoprecipitating HA-tagged small TbTims from the corresponding samples, we observed that TbTim9-HA and TbTim8/13-HA co-precipitated a fraction of TbTim10-Myc (60% of the input), but TbTim10-HA, TbTim12-HA, and TbTim13-HA precipitated less than 10% of TbTim10-Myc (Fig. 6B, lower panel, and D). This further indicates that interactions between TbTim10 with TbTim9 and TbTim10 with TbTim8/13 are stronger in comparison with other small TbTims (Fig. 6B-D). This is also correlated with our Y2H results. HA-antibody didn’t precipitate any protein from the mitochondrial extract that only have TbTim10-Myc. TbTim17 was co-precipitated from all samples similarly. VDAC was not precipitated by HA antibody from any samples, as expected. Due to some technical difficulties we failed to generate TbTim11-HA expressed *T. brucei* cell line, therefore, we couldn’t speculate the association of this proteins with other small TbTims. Together, our co-immunoprecipitation results showed that 1) TbTim10 has stronger association with TbTim9 and TbTim8/13 than other small TbTims, 2) TbTim10-Myc co-precipitated TbTim12-HA less in comparison to TbTim9-HA and TbTim8/13-HA, suggesting that TbTim9 and TbTim8/13 may form complexes either with TbTim10 or TbTim12, and 3) TbTim13 has stronger association for TbTim17 relative to TbTim10 with TbTim17. Therefore, we speculated that TbTim9 and TbTim8/13 are making a core complex either with TbTim10 or TbTim12. TbTim13 is strongly associated with the TbTIM17 complex, thus the loss of TbTim13 greatly hampered the integrity of this complex.

### Each of the small TbTims is a part of the smaller 70 kDa small TbTim complexes, except TbTim13, which is present in a larger complex (>800 kDa) and co-fractionates with TbTim17

Next, we analyzed the complexes formed by the small TbTims using size exclusion chromatography (SEC). Previously, we performed BN-PAGE of the digitonin-solubilized mitochondrial supernatant from *T. brucei* that expressed TbTim10-Myc and showed that TbTim10-Myc is present in smaller complexes (∼70 kDa, and ∼150 kDa) and in a larger complex (>800 kDa), which is similar in size to TbTIM17 complex (35). Here, we used *T. brucei* cells that expressed a pair of small TbTims with Myc and HA tags (TbTim10-Myc+TbTim9-HA, TbTim10-Myc+TbTim10-HA, TbTim10-Myc+TbTim8/13-HA, TbTim10-Myc+TbTim12-HA, and TbTim10-Myc+TbTim13-HA). Isolated mitochondria were solubilized with digitonin (1%) and soluble supernatants were fractionated on a gel-filtration column (SEC 650, 10 × 300). Each fractions (1–18) were analyzed by SDS-PAGE and immunoblot analysis using Myc-, HA-and TbTim17 antibodies. Results from fractions 1-13 (equivalent to vol 37-50 ml) are shown (Fig. 7A, C, E, G, I, and K). Other fractions (14–18) were negative to the antibodies tested. Results showed that TbTim10-Myc was enriched in fractions 3 and 6-7, whereas TbTim17 was enriched in fraction 2, when mitochondrial extract from TbTim10-Myc was analyzed (Fig. 7A and B). We calibrated the column with known molecular weight marker proteins and determine the sizes of the protein complexes in fractions 6 and 2 as ∼66 kDa, and 1964 kDa, respectively (Fig. S3). Therefore, we observed that TbTim10-Myc is present in a smaller complex (∼70 kDa) and in a larger complex closer in size with the TbTIM17 complex. When we expressed TbTim8/13-HA along with TbTim10-Myc and analyzed mitochondrial protein complexes by the same column, we found that both TbTim10 and TbTim8/13 were co-eluted in the fractions 6 and 7 (Fig. 7C and D). This is corelated with our co-immunoprecipitation results, where we showed that these two small TbTims were co-precipitated when pulled down by either Myc or HA antibodies (Fig. 6). A part of TbTim8/13-HA was detected in fraction 2 with TbTim17 (Fig. 7C and D). When we analyzed the mitochondrial extract from TbTim10-Myc+TbTim9-HA expressed *T. brucei*, we also found TbTim9-HA and TbTim10-Myc were found together in fractions 6 and 7 and a portion of these proteins were eluted with TbTim17 in fractions 2 and 3 (Fig. 7E and F). When TbTim12-HA was co-expressed with TbTim10-Myc, a larger fraction of TbTim12-HA was enriched in fractions 6 and 7 (Fig. 7G and H). However, in this sample a significant part of TbTim10-Myc was found in complexes with intermediate sizes between the larger and the 70 kDa complex (Fig. 7G and H). This suggests that in the presence of abundant TbTim12-HA, the 70 kDa complex is formed with TbTim12 and excess TbTim10-Myc associates with the other intermediate complexes. Co-expression of TbTim10-Myc with TbTim10-HA and analysis of the mitochondrial protein complexes by SEC 650, showed that TbTim10-HA behaved similarly with TbTim12-HA and primarily eluted with fractions 6 and 7; however, TbTim10-Myc levels were reduced in these fractions and found instead in the intermediate complexes (Fig. 7I and J). This suggests that due to expression of TbTim10 both as Myc and HA tags, some of the TbTim10-Myc and TbTim10-HA in the 70 kDa complex, and a part of these proteins was found in the intermediate complexes. In contrast to other small TbTims, TbTim13-HA was primarily enriched in fraction 2 along with TbTim17 and not found in fractions 6 or 7, indicating that TbTim13-HA does not form a smaller (70 kDa) complex (Fig. 7K and L). As mentioned before, in TbTim10-Myc+TbTim13-HA cells, levels of TbTim10-Myc were significantly reduced. Thus, a reduced level of TbTim10-Myc was eluted at fraction 2 (Fig. 7K and L). These results indicated that overexpression of TbTim13 reduced the assembly and stability of TbTim10. Immunoblot analysis of the column fractions using mHsp70 and VDAC antibodies showed that mHsp70 is eluted in a wide range of fractions (2–8) (Fig. 7M), which is not unexpected, since it is known that mHsp70 is associated with multiple mitochondrial complexes (40, 41). However, we noticed that mHsp70 was enriched in the fractions 6 and 7, which could be the monomeric form of this protein and this pattern of elution was consistent for all samples. Alternatively, VDAC was primarily eluted in fractions 2 and 3, which is likely the multimeric form of VDAC, as expected (Fig. 7M) and this pattern was consistent for all samples. Therefore, with these results along with our co-precipitation and RNAi data, we postulated that 1) TbTim13 is not present in a smaller (70 kDa) complex but rather associates only with the larger TbTIM17 complex, and it is crucial for the TbTIM17 complex assembly/stability. 2) TbTim10 and TbTim13 may compete for binding to TbTim17 as TbTim13 has stronger binding with TbTim17, thus overexpression of TbTim13 dissociates TbTim10 from the complex and reduces its stability. 3) TbTim9, TbTim10, TbTim8/13, and TbTim12 each forms the smaller (70 kDa) complex. 4) When TbTim10-Myc (inducible) and TbTim10-HA (constitutive) are expressed simultaneously, then TbTim10-HA mostly present in the 70 kDa complex and excess TbTim10-Myc is mostly present in the intermediate complexes, which correlated with our co-immunoprecipitation results that showed TbTim10-Myc and TbTim10-HA do not co-precipitated very well. From these results, we speculated a model for the small TbTim complexes (Fig. 8). TbTim9, TbTim10, TbTim8/13, and TbTim12 form multiple heterohexameric complexes, TbTim10 and TbTim12 could be present in different 70 kDa complexes, however TbTim13 stably associates with the TbTIM17 complex, and the small TbTims heterohexameric complexes dynamically associate with the larger TbTIM17 complex for protein translocation.

**Figure 7.**
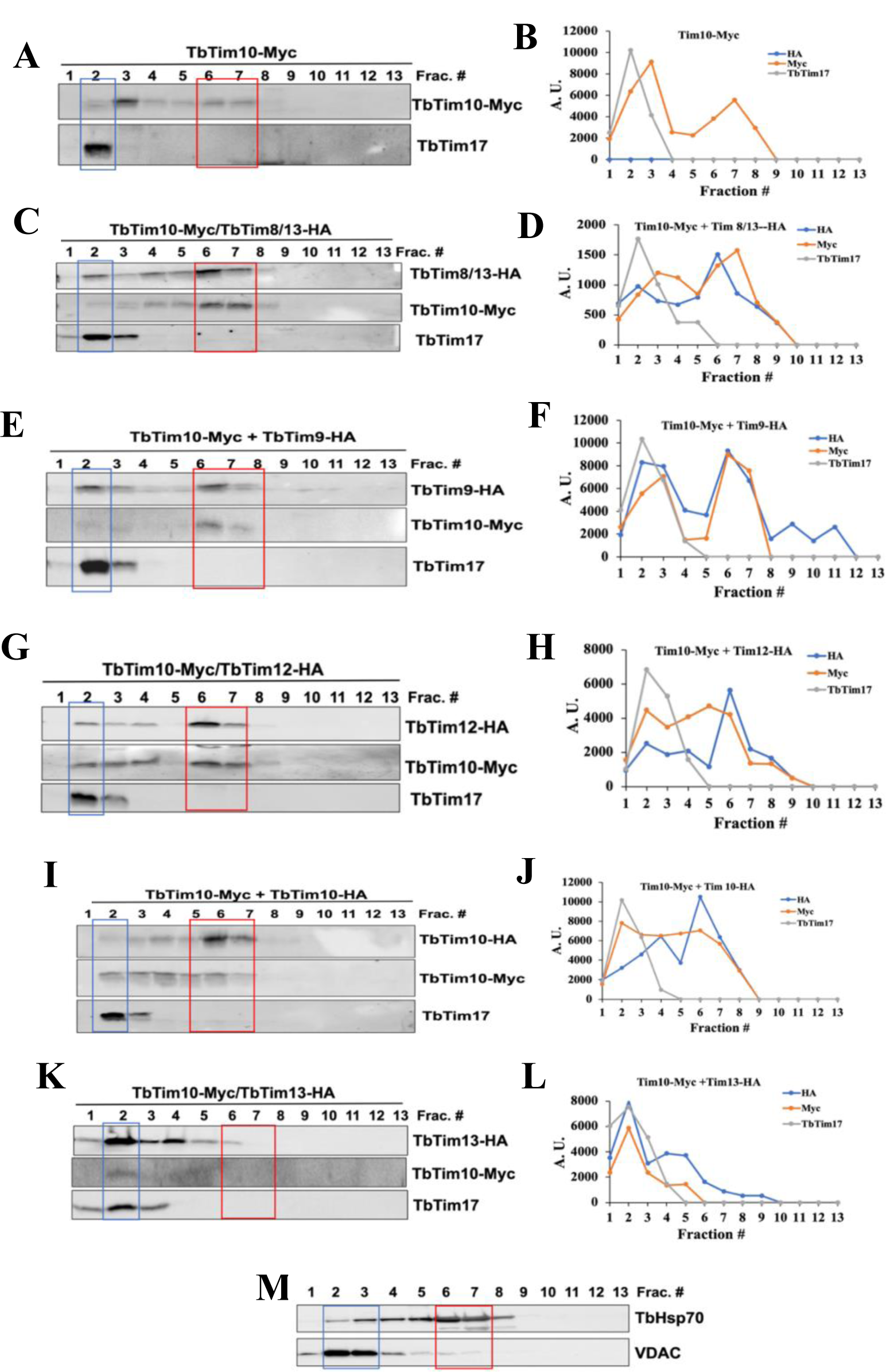
Analysis of the small TbTims complexes in *T. brucei*. Mitochondria isolated from the TbTim10-Myc, TbTim10-Myc/TbTim8/13-HA, TbTim10-Myc/TbTim9-HA, TbTim10-Myc/TbTim12-HA, TbTim10-Myc/TbTim10-HA, and TbTim10-Myc/TbTim13-HA *T. brucei* cells were solubilized with digitonin. The soluble supernatants were subjected to separate on SEC 650 column. Proteins were eluted with 1X Native buffer (20 mM Tris, pH 7.0, 50 mM NaCl, 1 mM EDTA, 10% glycerol, 1 mM PMSF, and 0.1% digitonin). (A, C, E, G, I, and K) Fractions (1–13) were analyzed by SDS-PAGE and immunoblot analysis using HA and Myc antibodies. (M) mHSP70 and VDAC were used as standard mitochondrial proteins. Fractions 2 and 6-7 are in blue and red boxes. (B, D, F, H, J, and L) Myc-and HA-tagged small TbTims and TbTim17 bands were quantitated by ImageJ software for each run and plotted against the fraction numbers in Excel. Band intensity was presented in arbitrary units (A.U.). Runs from multiple experiments from each sample were reproducible.

**Figure 8.**
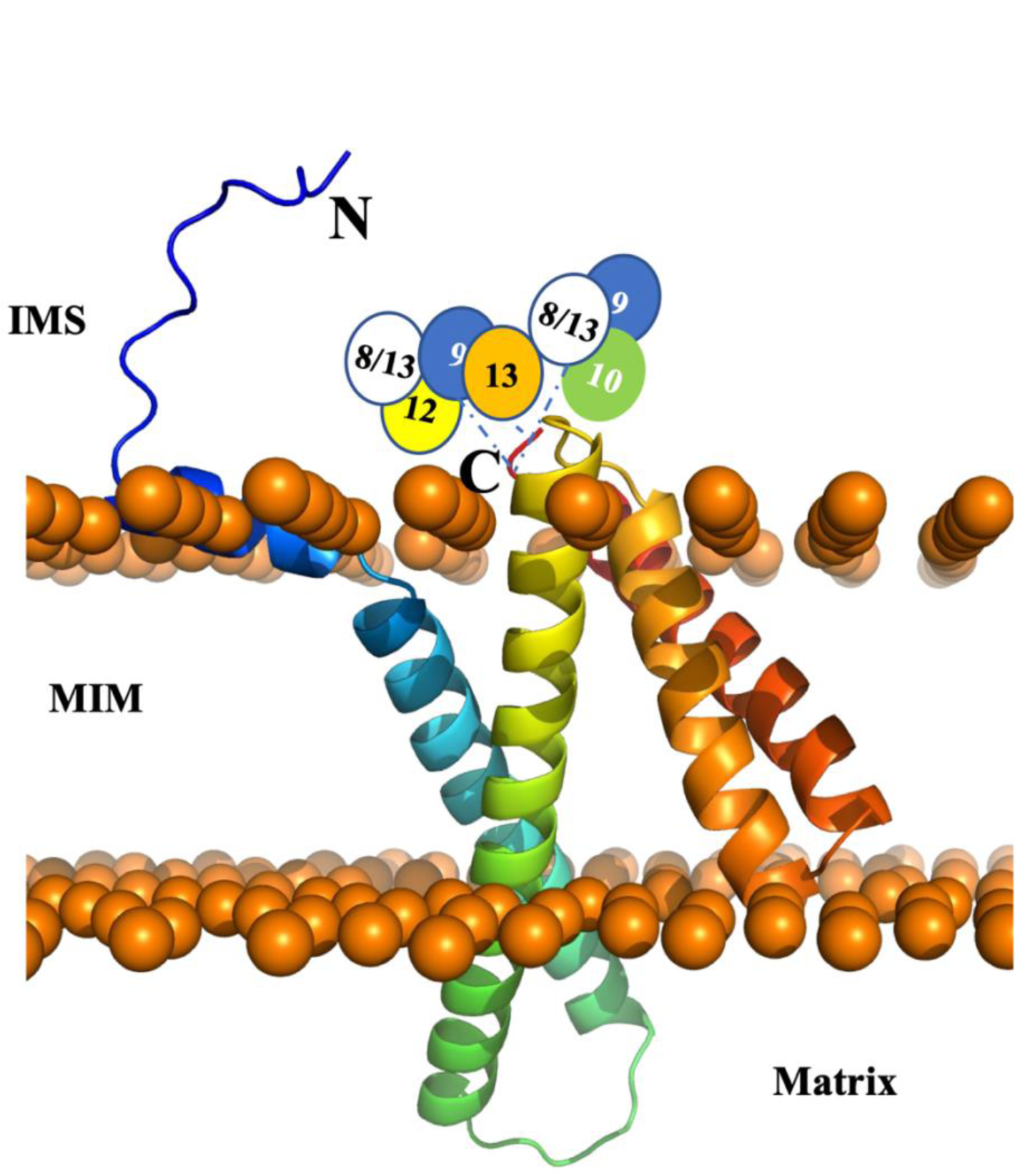
Model for the small TbTims complexes with TbTim17. The TbTim17 model was generated using AlphaFold within the ChimeraX software. The model was then uploaded to CHARMM-gul server and oriented within a mitochondrial membrane through the OPM server. Using the CHARMM-gul server, the model was assembled and equilibrated. The N-and C-terimini are marked. Mitochondrial inner membrane (MIM), intermembrane space (IMS), and matrix are labeled. The putative small TbTim complexes as for example (TbTim9-TbTim8/13-TbTim10) and (TbTim9-TbTim8/13-TbTim12) are placed near the C-terminal of TbTim17 and linked by dashed blue lines. TbTim13 stays attached to the C-terminal of TbTim17.

## Discussion

The TbTIM17 complex is unique in *T. brucei* in multiple aspects. It consists of several trypanosome-specific proteins, and it is involved in import of both the N-terminal MTS-containing and the internal targeting signal-containing hydrophobic MIM proteins as well as tRNAs (42, 43). It is speculated that TbTim17 performs these complex functions by dynamically associating with different components; however, a complete picture is far from clear. *T. brucei* possesses six essential, small TbTims, which were discovered as parts of these complex; however, there was limited information regarding their interaction pattern among themselves and with TbTim17. Here, using Y2H analysis and co-immunoprecipitation studies, we showed that TbTim8/13 has stronger interactions with TbTim9 and TbTim10 in comparison to others, whereas TbTim13 showed stronger association with TbTim17. This observation was also supported by analysis of the small TbTim complexes using SEC, which revealed that TbTim9, TbTim8/13, TbTim10, and TbTim12 capable to form 70 kDa complexes, which could be similar to the heterohexameric complexes of the small Tims found in yeast and mammals (22, 23), whereas TbTim13 stays associated with the larger TbTIM17 complex and is crucial for its integrity. In addition, we found that the C-terminal domain of TbTim17 is likely the point of contact with small TbTims. Therefore, it appears that TbTim13 may act as ScTim12 or HsTim10b and associate as a stable part of the larger translocase complex, while the other small TbTims forms the characteristic 70 kDa complex in different combinations and dynamically associate with the larger TbTIM17 complex. Together, these studies increased our understanding about the single TIM complex in *T. brucei*.

After comparing the effect of knockdown of each of the small TbTims in parallel, we found that the loss of TbTim13 has more drastic effect on the levels of the TbTIM17 complex as well as TbTim17 steady-state protein levels in comparison to the knockdown of other small TbTims. TbTim10 and TbTim8/13 also showed a similar reduction of TbTIM17 complex, but it takes a longer induction of RNAi. Wenger et al., studied TbTim11, TbTim12, and TbTim13 and reported that TbTim13 is most important to maintain TbTIM17 complex levels, but not for TbTim17 protein (34). However, we found a significant loss of TbTim17 protein due to TbTim13 RNAi. This discrepancy could be due to differential extent of knockdown or different laboratory strain of *T. brucei*. A significant loss of the steady-state levels of TbTim17 in our experiment is possibly because the unassembled TbTim17 was degraded in the absence of TbTim13. In fact, by quantitative proteomics analysis, we found a 5-fold increase of the Clp protease, a quality control protease of mitochondria, due to knockdown of TbTim13 (Supplemental file S1, Fig. 5A). In any case, it would be interesting to elucidate the exact role of TbTim13 in TbTim17 biogenesis/stability.

From our proteomics analysis, we found that depletion of any of these small TbTims creates a stress response phenomenon, which is indicated by upregulation of multiple heat shock proteins, chaperones, and mitochondrial proteases. This could be a direct effect of small TbTims knockdown or due to a reduction of TbTIM17 complex in small TbTims RNAi cells. The latter is more likely the cause, as we observed that upregulation of these stress-related proteins is maximum for TbTim13 RNAi that causes maximum adverse effect on the TbTIM17 complex (Supplementary file S1). Interestingly, we found significant downregulation of Atom46, a receptor of the ATOM translocase, particularly in TbTim9, TbTim11, and TbTim8/13 RNAi cells. It is possible that in the absence of these small TbTims, Atom46 is downregulated to control the rate of mitochondrial import of incoming proteins. Alternatively, these small TbTims could have specific roles on biogenesis or stability of this MOM protein. TbTim42, a component of the TbTIM17 complex, was downregulated in most of the small TbTim RNAi mitochondria, but less for TbTim13 knockdown in comparison to others. This is an interesting phenomenon that needs further investigation. It is known that MCPs are the substrate for small Tims in other eukaryotes. This could also be the case for *T. brucei*, as this parasite possesses a large group of MCPs. We found two such proteins, MCP9 and MCP12, were significantly downregulated due to small TbTims knockdown, indicating that these are involved in the biogenesis of MCPs. Most of the other MCPs may have higher abundance or higher half-life, thus couldn’t be detected among the downregulated proteins. Several COX subunits, particularly COV were reduced due to depletion of any of the small TbTims. It is known that human Tim8 and Tim13 plays role in COX assembly; however, it requires further investigation if small TbTims have any such roles.

Analysis of the small TbTim complexes by SEC analysis showed that TbTim10 eluted together with TbTim9 or TbTim8/13 in fractions 6-7, which indicates that they could form the heterohexameric complexes. The stoichiometry of the subunits in the 70 kDa complex needs to be further determined; however, we can assume that similar to its counterpart in yeast and human, it could be the (TbTim10)_3_-(TbTim9)_3_, (TbTim10)_3_-(TbTim8/13)_3_, or (TbTim8/13)_3_-(TbTim9)_3_ complexes. We also cannot exclude the possibility that all three proteins are present in the same complexes, i.e. (TbTim8/13)_3_-(TbTim9)_2_-TbTim10 or (TbTim8/13)_2_-(TbTim9)_3_-TbTim10. We observed that TbTim10-HA didn’t coprecipitate well TbTim10-Myc. We also observed that TbTim10-HA is primarily present in the 70 kDa complex, whereas TbTim10-Myc is mostly found in multiple intermediate complexes. This indicates that the constitutively expressed TbTim10-HA saturates the 70 kDa complex, therefore TbTim10-Myc expressed in excess after induction was not assembled in the smaller complex. TbTim12-HA and TbTim10-Myc were present in the 70 kDa complex when co-expressed, however excess TbTim10-Myc was also found in the larger intermediate complexes, suggesting that TbTim12 is not present in the same 70 kDa complex with TbTim10-Myc but may have some common constituents. We observed that TbTim10 and TbTim13 didn’t co-precipitate from mitochondrial extract, however, TbTim13 strongly co-precipitate TbTim17. Furthermore, we demonstrated TbTim13 co-eluted with TbTim17 in fractions 2-3 by SEC and not in fractions 6-7. Therefore, we conclude that TbTim13 associates with the TbTIM17 complex and not present in the 70 kDa complex.

The structure of the human TIM22 complex have recently been resolved by Cryo-EM at a resolution of 3.7 Å (39). This structure revealed that the N-terminal helices of HsTim22 interacts with 2 heterohexameric small Tim complexes, Tim9_3_-Tim10_3_ and Tim9_2_-Tim10a_3_-Tim10b. The other components of the HsTIM22 complex, like acylglycerol kinase and Tim29 also interact with the small Tim complexes. Currently it is not clear the stoichiometric ratios of the components of the TbTIM17 complex. From the abundance of the TbTim17 in the TbTIM complex, it suggests that more than one copy of TbTim17 is present per complex. We found that the C-terminus of the TbTim17 interacted with all the small TbTims stronger than the N-terminus, in Y2H analysis, suggesting that the C-terminal end is likely to hold the small TbTim complexes. However, whether or not this is true in *T. brucei* mitochondria requires further verification. Overall, our studies have advanced the knowledge about the interaction of the small TbTims with each other and with TbTim17, as well as how they form different complexes. Future studies by purification of these complexes and Cryo-EM analysis will uncover more details about the structure of this unique translocase complex in *T. brucei*.

### Experimental procedures

#### Cell maintenance, growth medium, and cell growth analysis

The procyclic form of the *T. brucei* 427 doubly resistant cell line (29–13) expressing a tetracycline repressor gene and a T7 RNA polymerase was grown in SDM-79 medium supplemented with 10% fetal bovine serum, G418 (15 µg/ml), and hygromycin (50 µg/ml) (44). To measure growth, cells were seeded at a cell density of 3 x 10^6^ cells/ml in fresh medium containing the appropriate antibiotics. Cell numbers were counted at different time points (0-8 days) using a Neubauer hemocytometer. The log of cumulative cell numbers was plotted versus time (in days) of incubation.

#### Generation of plasmid constructs and *T. brucei* transgenic cell lines

*T. brucei* cell line that express TbTim10-Myc upon induction with doxycycline was developed previously (35). For generation of double-tagged small TbTim cell lines (TbTim10-Myc/TbTim9-HA, TbTim10-Myc/TbTim10-HA, TbTim10-Myc/TbTim12-HA, TbTim10-Myc/TbTim13-HA, and TbTim10-Myc/TbTim8/13-HA), the open reading frames (ORFs) of TbTim9, TbTim10, TbTim12, TbTim13, TbTim8/13 was PCR-amplified using *T. brucei* 427 genomic DNA as the template and the corresponding sequence-specific primers (Table S1). The forward and reverse primers were designed with addition of restriction sites for HindIII and XbaI at the 5’ ends, respectively. The PCR products were cloned into the modified pHD1344 vector, where the coding sequence was inserted within the HindIII and XbaI restriction sites. Generated constructs were linearized by NotI digestion and transfected into *T. brucei* TbTim10-Myc cell line as described (35). Transfected cells were selected with puromycin (1.0 µg/ml). The TbTim9 RNAi, TbTim10 RNAi, and TbTim8/13 RNAi cells were developed previously (35). The constructs for TbTim11 RNAi, TbTim12 RNAi, and TbTim13 RNAi were generated by PCR amplification of the ORFs of TbTim11, TbTim12, and TbTim13, using *T. brucei* genomic DNA as the template and sequence-specific primers (Table S1). The forward and reverse primers were designed with addition of restriction sites for HindIII and BamHI at the 5’ ends, respectively. The PCR products were cloned into tetracycline-inducible p2T7^Ti^-177 RNAi vector between the HindIII and BamHI restriction sites (45) Plasmid DNA was linearized by NotI digestion and transfected into *T. brucei* 29-13 cells. Transfected cells were selected with puromycin (1.0 µg/ml).

#### RNA isolation and quantitative RT-PCR analysis

RNA was isolated from *T. brucei* cells using RNeasy miniprep isolation kit (Qiagen) and digested with amplification grade DNase (1U/µl) for 1 h before first-strand cDNA synthesis, which was performed using an iScript cDNA synthesis kit (Bio-Rad). The resulting cDNA was amplified using specific primers designed from the 3’-untranslated region sequence of the target genes for small TbTims (Table S1) to detect endogenous TbTim11, TbTim12, and TbTim13 mRNA levels but not the double stranded RNA generated from the p2T7^TI^-177 RNAi construct. Primers for amplification of tubulin cDNA were generated from the ORF of the tubulin gene (Table S1).

#### Yeast two-hybrid analysis

The ORFs of TbTim11, TbTim12, and TbTim13 were subcloned into yeast expression vectors pGADT7 and pGBKT7 (Takara) to generate the bait and prey plasmids (46). TbTim9, TbTim10, and TbTim8/13 were cloned previously in the pGADT7 and pGBKT7 vectors (35). Similarly, the two helices of each small TbTims were cloned in the same vectors. The coding regions of TbTim17 N-terminal (1-30 aa), loop2 (140-152 AAs) and C-terminal (93-107 AAs) as well as the ORF of the TbTim17 full-length protein were subcloned into yeast expression vectors pGADT7 and pGBKT7. Approximately 2 µg of each of the bait and prey plasmids in different combination pairs was co-transformed into the *Saccharomyces cerevisiae* Y2H Gold strain (Takara) using the lithium acetate method (35). Co-transformed yeast cells were plated on SD medium lacking leucine (-leu) and *tryptophan (-trp)* and allow to grow for 3 days at 30°C. Yeast clones that grew were then plated on SD –leu/–trp/–his medium that was lacking leucine, tryptophan, and histidine to select for protein-protein interactions. SD –leu/–trp/–his plates were also supplemented with 2.0, 3.5, and 5.0 mM 3-amino-1,2,4-triazole (AT), which inhibits Y2H yeast cell growth due to leaky expression of the *HIS3* gene (46–48), to limit the occurrence of false positives. Inoculated plates were allowed to grow at 30°C for 3 to 5 days. To confirm positive readouts, this process was repeated at least three times with individual clones.

#### Sub-cellular fractionation and crude mitochondria isolation

Fractionation of *T. brucei* procyclic form cells was performed as described (35). Briefly, 2×10^8^ cells were pelleted and re-suspended in 500 μL of SMEP buffer (250 mM sucrose, 20 mM MOPS/KOH, pH 7.4, 2 mM EDTA, 1 mM PMSF) containing 0.03% digitonin and incubated on ice for 5 min. The cell suspension was then centrifuged for 5 min at 6,800 × g at 4°C. The resultant pellet was considered as the crude mitochondrial fraction, and the supernatant contained soluble cytosolic proteins. Mitochondria were isolated from the parasite after lysis by nitrogen cavitation in isotonic buffer (36). The isolated mitochondria were stored at a protein concentration of 10 mg/ml in SME buffer containing 50% glycerol at −70°C. Before use, mitochondria were washed twice with nine volumes of SME buffer to remove glycerol (36).

#### SDS-PAGE and immunoblot analysis

Proteins from whole-cell lysates or cytosolic or mitochondrial extracts were separated on a 15% SDS polyacrylamide gel, transferred to a nitrocellulose membrane, and immunodecorated with polyclonal antibodies for TbTim17 (Tb927.11.13920) (49), TbAAC (Tb927.10.14820) (49), VDAC (Tb927.2.2510) (50), TbPP5 (Tb927.10.13670) (51), and mtHsp70 (Tb927.6.3740) (52), and *T. brucei*-tubulin (Tb927.1.2350) (54). Anti-Myc and anti-HA polyclonal antibodies were purchased from commercial sources (Abcam and Thermo-Fisher, respectively). Blots were developed with appropriate secondary antibodies and an enhanced chemiluminescence kit (Thermo-Fisher).

#### BN-PAGE analysis

Mitochondrial proteins (200 µg) were solubilized in 100 µl of ice-cold 1x native buffer (Thermo-Fisher) containing 1% digitonin. The solubilized mitochondrial proteins were clarified by centrifugation at 100,000 *g* for 30 min at 4°C. The supernatants were mixed with G250 sample additive (Invitrogen) and were electrophoresed on a precast (4% to 16%) bis-Tris polyacrylamide gel (Thermo-Fisher), according to the manufacturer’s protocol. Protein complexes were detected by immunoblot analysis. Molecular size marker proteins apoferritin dimer (886 kDa) and apoferritin monomer (443 kDa), amylase (200 kDa), alcohol dehydrogenase (150 kDa), and bovine serum albumin (66 kDa) were electrophoresed on the same gel and visualized by Coomassie staining. Proteins were transferred to a nitrocellulose membrane for Western blot analysis.

#### Co-immunoprecipitation assay

Mitochondria (600 μg) were solubilized in 300 µl of 1 x cold native buffer (50 mM [Tris pH 7.2], 50 mM NaCl, 10% [wt/vol] glycerol, 1 mM PMSF, 1% digitonin) and incubated on ice for 1 h. The solubilized mitochondria were centrifuged at 100,000 *g* for 30 min. An aliquot (50 μl) of the supernatant was mixed with 50 µl of 2 x Laemmli sample buffer and served as the input sample. The remaining supernatant (∼250 µl) was mixed with 25 µl of anti-myc or anti-HA conjugated agarose bead slurry (Sigma-Aldrich) and allowed to incubate for 12 h at 4°C with constant gentle inversion. The beads were separated by centrifugation at 1,000 g for 1 min. An aliquot (50 μl) of the supernatant was mixed with 50 µl of 2 x Laemmli sample buffer and served as the unbound fraction. The beads were then washed three times in wash buffer (50 mM Tris [pH 7.2], 50 mM NaCl, 10% [wt/vol] glycerol, 1 mM PMSF, 0.1% digitonin) to remove nonspecifically bound proteins. The washed beads with bound proteins were resuspended in 50 µl of 1 x Laemmli sample buffer and served as the bound fraction. The samples were then separated on a 15% SDS polyacrylamide gel and transferred to a nitrocellulose membrane. Blots were developed with appropriate secondary antibodies and an enhanced chemiluminescence kit (Thermo-Fisher).

#### Size exclusion chromatography

Mitochondrial proteins (4 mg) were solubilized in 1.25 ml of 1X Native buffer (20 mM Tris, pH 7.0, 50 mM NaCl, 1 mM EDTA, 10% glycerol, 1 mM PMSF, and 1% digitonin). The solubilized proteins were loaded on a ENrich^TM^ SEC 650 10 × 300 column (BioRad) and run on an NGC system (BioRad). The Proteins were eluted with the same buffer except the digitonin concentration was reduced to 0.1%. After discarding the void volume (6.0 ml) 18 fractions (1.0 ml each) were collected. Proteins in each fraction were analyzed by SDS-PAGE and immunoblotting using antibodies for Myc, HA, TbTim17, VDAC, and mHsp70. Gel filtration molecular weight markers were run separately on the column under same conditions and were detected in the eluted fractions by SDS-PAGE and CB-staining (Supplemental Fig. S3). The sizes the small TbTims complexes and TbTim17 complexes were calculated from a standard curve in which the log10 values of molecular sizes of marker proteins were plotted against the elution volumes.

## Mass spectrometry analysis

For shotgun mass spectrometry analysis of the mitochondria samples from the parental control and each of the small TbTim RNAi cells (day 4 post-induction), equal amounts of proteins were solubilized in 2x SDS-PAGE sample buffer and run on a denaturing gel. After the proteins had penetrated 1 cm into the resolving gel, it was stained with CB and the stained area was excised for further analysis by trypsin digestion and MS/MS analysis as described (40).

### Bioinformatics analysis

Bioinformatics pathway enrichment analysis was performed using the database for Annotation, Visualization and Integrated Discovery Bioinformatics resources (DAVID version 6.7) (55), the Search Tool for the Retrieval of Interacting Genes/Proteins (STRING version 11.0) (56), and the Kyoto Encyclopedia of Genes and Genomes (KEGG) (57) databases and search tools. The identified enriched pathways with *P* values of <0.05 were selected. Significantly enriched pathways were determined using a significance threshold of an FDR corrected P-value <0.005. Protein family enrichment was done using the STRING database with Protein families (Pfam) as the reference database (58). Functional domain analysis of proteins was performed using InterPro version 84.0 to classify protein families and predicted active domains (59).

## Author Contribution

M. C., L. Q. G., F. S. G., and C. D. conceptualization and execution, V. P. and A. C. proteomics and bioinformatics analysis, M. K., A. T., and J. T. S. generation of clones, J. D. SEC analysis, M. C., and L.Q.G. writing-original draft, all authors reviewing and editing.

## Funding and additional information

This work was supported by NIH grant 1RO1AI125662 (M.C.). L. Q. G. was supported by 2R25GM059994 from NIGMS. This work was accomplished in part through the use of the Meharry Medical College Core Facilities, which are supported by NIH Grants MD007586, CA163069, and S10RR025497. The content is solely the responsibility of the authors and does not necessarily represent the official views of the National Institute of Health.

## Abbreviations

The abbreviations used are: AA, amino acid; AAC, ADP/ATP carrier; ATOM, archaic translocase of outer mitochondrial membrane; AD, activation domain; AT, 3-amino-1,2,4-triazole; BD, binding domain; BN, blue-native; CB, Coomassie blue; COX, cytochrome oxidase; DDO, double dropout; EM, electron microscope; HA, hemagglutinin; HRP, horseradish peroxidase; IMS, intermembrane space; MCP, mitochondrial carrier protein; MIM, mitochondrial inner membrane; MOM, mitochondrial outer membrane; MudPIT, multidimensional protein identification, ORF, open reading frame, PAM, presequence translocase associated motor; PMSF, phenyl methyl sulfonyl fluoride; SEC, size exclusion chromatography; SME, sucrose 3-N-morpholino propanesulfonic acid (MOPS) ethylenediaminetetraacetic acid; TDO, triple dropout; TIM, translocase of the inner mitochondrial membrane; VDAC, voltage-dependent anion channel; Y2H, yeast two hybrid.

