## Supplemental Table-1, supplemental figure S1, S2, and S3 for "Unique interactions and functions of the mitochondrial small Tims in *Trypanosoma brucei*"

**Table S1**

**Primer Name**	**DNA Sequence (5’-3’)**

TbTim9-HA Forward	GATCAAGCTTATGCGCCTGGCTG

TbTim9-HA Reverse	GATCTCTAGACAACTTCAACATTTG

TbTim10-HA Forward	GATCAAGCTTATGCAGCCACCTC

TbTim10-HA Reverse	GATCTCTAGATTCGTTCTCCGAC

TbTim8/13-HA Forward	GATCAAGCTTATGAATCAGTCTAGTTC

TbTim8/13-HA Reverse	GATCTCTAGACCTTTCCTTTGCTTG

TbTim12-HA Forward	GATCAAGCTTATGGGTCAGGACCAATCC

TbTim12-HA Reverse	GATCTCTAGAACGACTAAGTTCCTTTC

TbTim13-HA Forward	GATCAAGCTTATGCAACCCCCAACCCCAC

TbTim13-HA Reverse	GATCTCTAGAGACCCCTCCCCCCTG

TbTim11 RNAi Forward	GATCAAGCTTATGCAGAGCCAAATGATGC

TbTim11 RNAi Reverse	GATCGGATCCCTACTCCTCCATACCGAGGGATC

TbTim12 RNAi Forward	GATCAAGCTTATGGGTCAGGACCAATCC

TbTim12 RNAi Reverse	GATCGGATCCTTAACGACTAAGTTCCTTTC

TbTim13 RNAi Forward	GATCAAGCTTATGCAACCCCCAACC

TbTim13 RNAi Reverse	GATCGGATCCCTAGACCCCTCCCCCCTG

TbTim11-qRT Forward	TTTTGCCGGTCAAACAGAGC

TbTim11-qRT Reverse	CCAGCCACAAGTGATTGAAGTG

TbTim12-qRT Forward	ATTACCCTTCGGCGTCAAGG

TbTim12-qRT Reverse	GCTTTCACCCAGTCTCCGAA

TbTim13-qRT Forward	GGCGAACTGGCAGTGATAGT

TbTim13-qRT Reverse	CGCTACTCTTGCAACCCTCA

*Restriction enzyme sites are underlined

A

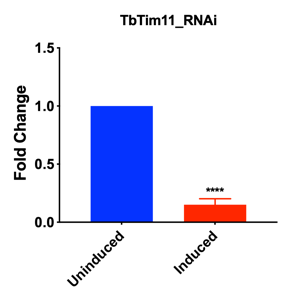

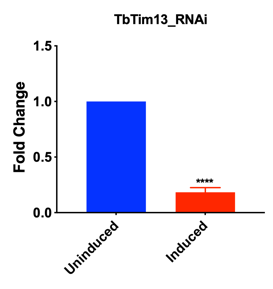

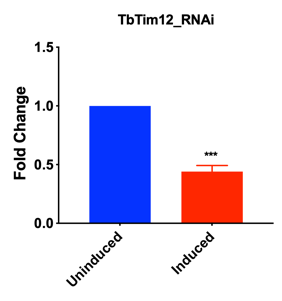

B

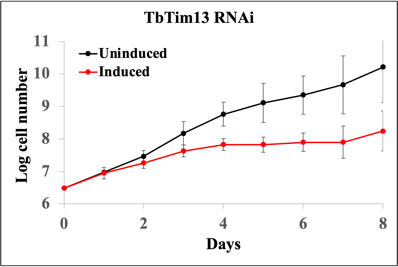

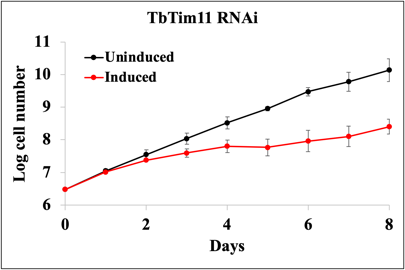

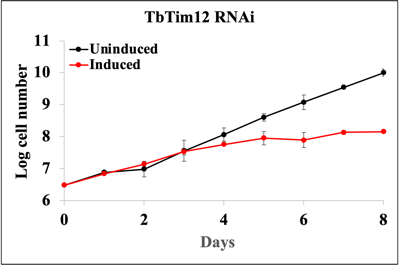

**Figure S1. Effect of small TbTim RNAi on the levels of specific transcripts and cell groeth in *T. brucei*.** A) qRT-PCR analysis of TbTim11, TbTim12, TbTim13 mRNA levels in TbTim11 RNAi, TbTim12 RNAi, TbTim13 RNAi *T. brucei* cells grown in the absence (Uninduced) and presence of doxycycline (1.0 μg/ml) for 48 h. The mRNA levels of the target transcript in the uninduced cells were considered as 100%. All transcript levels were normalized with the levels of the tubulin mRNA in each sample. Values were calculated from three independent replicates and are represented as scatterplots with error bars for standard deviations. Pvalues are 0.0001 (****) and 0.0004 (***). B) Cell growth analysis of TbTim11, TbTim12, and TbTim13 RNAi parasite in the absence (Uninduced) and presence (Induced) of doxycycline (1.0 μg/ml) for 8 days. The log of the cumulative cell numbers was plotted versus days post-induction. Standard errors were calculated from three independent experiment.

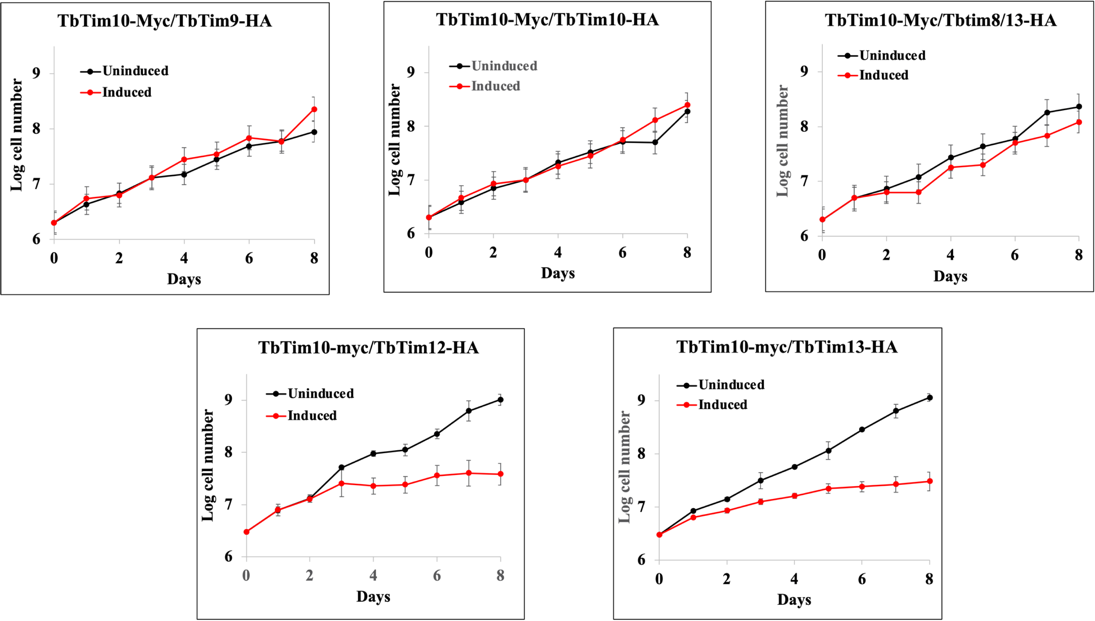

**Figure S2. Effect of overexpression of small TbTims on cell growth.** Cell growth analysis of T. brucei expressed TbTim10-Myc+TbTim9-HA, TbTim10-Myc+TbTim10-HA, TbTim10-Myc+TbTim8/13-HA, TbTim10-Myc+TbTim12-HA, and TbTim10-Myc+TbTim13-HA parasite in the absence (Uninduced) and presence (Induced) of doxycycline (1.0 μg/ml) for 8 days. The log of the cumulative cell numbers was plotted versus days post-induction.

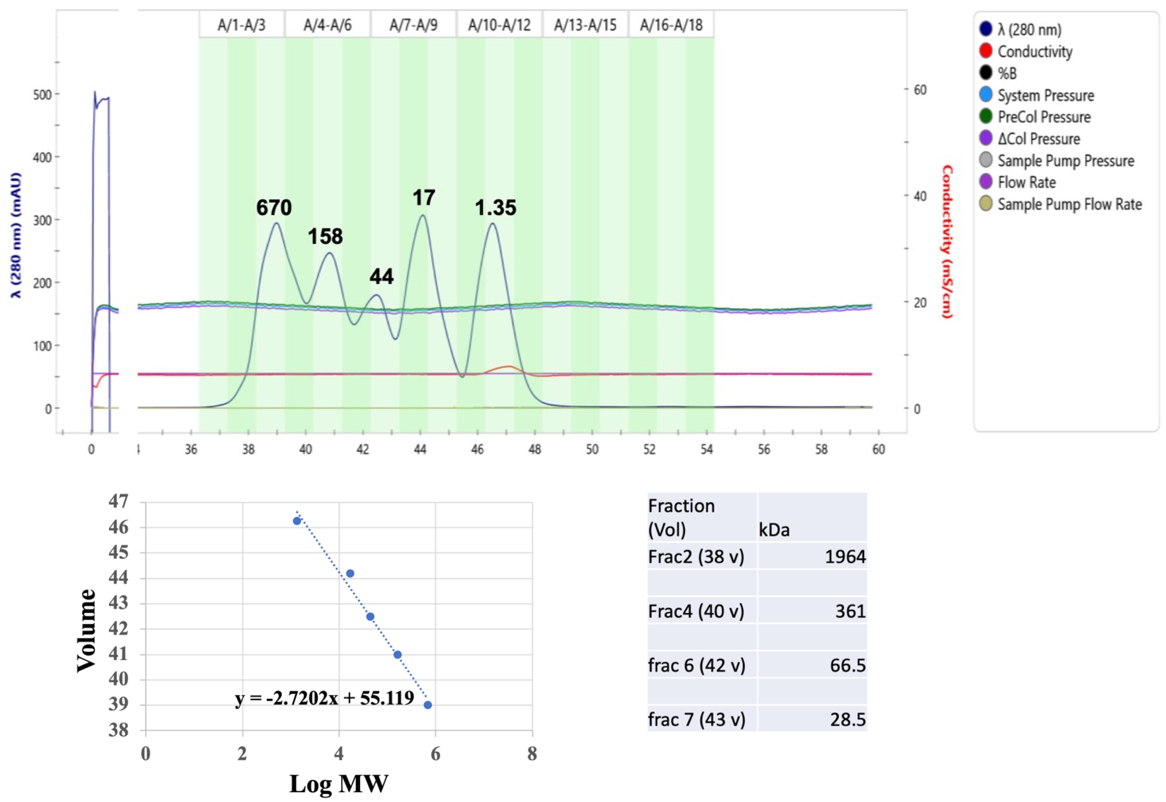

**Figure S3. Standard protein run on SEC650.** Standard molecular marker protein mixture (BioRad) (200 µl) was loaded on a ENrich^TM^ SEC 650 10 X 300 column (BioRad) and run on an NGC system (BioRad). Proteins were eluted with 1X Native buffer (20 mM Tris, pH 7.0, 50 mM NaCl, 1 mM EDTA, 10% glycerol, 1 mM PMSF, and 0.1% digitonin). Peaks for known molecular weight proteins; Thyroglobulin, 670 kDa; 𝛾-globulin, 158 kDa; Ovalbumin, 44 kDa; Myoglobin 17 kDa; and vitamin B12, 1.35 kDa are marked. Proteins were identified by color and by expected molecular sizes on a denaturing gel. A standard curve was generated by plotting elution volumes of the peaks versus log of molecular weight of the standard proteins. Unknown molecular weight of the mitochondrial protein complexes eluted at different volumes were calculated and shown in the table.
